# Tar Spot Disease Severity Influences Phyllosphere-Associated Bacterial and Fungal Microbiomes

**DOI:** 10.1101/2024.01.12.565617

**Authors:** Raksha Singh, Charles F. Crane, Sujoung Shim, Darcy E. P. Telenko, Stephen B. Goodwin

## Abstract

Tar spot, caused by the obligate fungal pathogen *Phyllachora maydis*, is a foliar disease of corn that has become a major economic concern in the United States. To test the hypothesis that *P. maydis* can interact with other foliar microorganisms, we investigated phyllosphere microbiomes in relation to corn inbreds with differential tar spot symptoms under natural infestation in the field. Leaf samples from sixteen inbred lines were assessed for tar spot symptoms, and bacterial and fungal microbiomes were characterized by paired-end sequencing on the Illumina MiSeq platform. Comparison of the phyllosphere microbiomes revealed distinct bacterial and fungal communities between resistant and susceptible lines. Bacterial and fungal species richness was significantly higher in resistant compared to susceptible inbred lines in a sample-specific manner. In contrast, there were no clear differences in diversity when including evenness of bacterial communities between the resistant and susceptible lines. Diversity of fungal communities differed significantly, particularly between twelve of the fourteen susceptible lines versus resistant lines. Plant-beneficial bacterial genera such as *Methylorubrum* and *Quadrisphaera* were associated with resistant lines, while *Pantoea, Deinococcus* and *Pseudomonas* were the least abundant. The second-most commonly detected fungus likely was a *Coniothyrium*, but whether it is the same species that was identified as a possible mycoparasite of *P. maydis* in Central and South America is not known. Fungal genera *Cladosporium, Papiliotrema, Cryptococcus, Tilletiopsis* and *Alternaria* were associated with resistant lines while *Sphaerellopsis* was the least-abundant genus. In contrast, *Puccinia, Sphaerellopsis* and *Phyllachora* were the dominant fungal genera in susceptible lines. Our findings imply that *P. maydis* infection may result in a distinct microbiota with lower diversity. Further analyses of these distinct microbiota between resistant and susceptible lines could lead to a better understanding of the potential role of foliar microbiomes in causing or resisting *P. maydis* infection.

---

Corn (maize; *Zea mays* subsp. *mays*) is one of the most widely grown crops in the world and is used for a wide range of food, feed and industrial applications. Among numerous risks to worldwide corn production, disease-related yield losses pose a serious threat to our capacity to attain long-term food and nutritional security, and inflict severe financial stresses to growers (Savary et al. 2019). Several foliar diseases, including northern leaf blight (NLB) caused by *Exserohilum turcium* (Hooker 1976)), southern leaf blight (SLB) caused by *Bipolaris maydis* (anamorph: *Cochliobolus heterostrophus*) (Ullstrup 1972), gray leaf spot caused by *Cercospora zeae-maydis* and *C. zeina* (Tehon and Daniels 1925; Crous et al., 2006), and common and southern rust caused by *Puccinia sorghi* and *P. polysora* (Cummins 1941), respectively, can contribute to yield loss.

Another foliar disease, tar spot, has recently emerged as a major threat to corn production, causing yield losses of up to 25 to 50% on susceptible hybrids under favorable conditions (Rocco da Silva et al. 2021; Valle-Torres et al. 2020). Tar spot is caused by the obligate biotrophic fungal pathogen *Phyllachora maydis* Maubl and was first identified in Indiana and Illinois in 2015 (Bajet et al. 1994; Bissonnette 2015; Ruhl et al. 2016). After infection, *P. maydis* produces black, glossy, elevated, round or oval stromata on the leaves and stems of corn. In some cases, a fish-eye symptom occurs when stromata are encircled by a tan-colored necrotic halo (Bajet et al. 1994; Hock et al. 1995; McCoy et al. 2019). Although *P. maydis* can infect corn at any stage of growth, the most severe symptoms develop during the silk (R1) and later stages.

Possible control strategies for tar spot include host resistance, biological control, and fungicidal chemicals. Host resistance should be the most effective, economical and environmentally benign strategy (Rocco da Silva et al. 2021; Valle-Torres et al. 2020) but is not widely available in commercial cultivars. A diverse panel of tropical and subtropical corn germplasm has been screened for tar spot resistance or tolerance with the intent to develop improved varieties (Cao et al. 2017; Lipps et al. 2022; Ren et al. 2022; Singh et al. 2022), but less is known about resistance in North American corn cultivars (Lipps et al. 2022; Singh et al. 2022). Biological control is also being investigated, and several taxa in the corn foliar microbiome have been found to inhibit specific pathogens (reviewed in Singh and Goodwin 2022). Thoroughly understanding the microbial response to *P. maydis* infection might reveal microbes associated with resistance that can suppress tar spot disease.

The phyllosphere contains diverse epiphytic and endophytic organisms that can exhibit commensal, parasitic, or mutualistic interactions with leaf tissue. Phyllospheric organisms can significantly affect their host and the ecosystem at large (Lindow and Brandl 2003). Disease resistance and susceptibility have been linked to the phyllospheric microbiome in a variety of species, including tomato (*Solanum lycopersicum*), poplar (*Populus*) , wheat (*Triticum aestivum L*.), rice (*Oryza sativa*), corn (*Zea mays*), and *Arabidopsis thaliana*, and disease can be altered by manipulating phyllospheric microbes (Berg & Koskella 2018; Busby et al. 2016; De Costa et al. 2006a, b; Massart et al. 2015; Ritpitakphong et al. 2016). Despite the importance of phyllospheric microorganisms to plant health, relatively little is known about their response to fungal foliar diseases. Only a few studies have probed the dynamics of microbial populations in the corn phyllosphere in response to fungal pathogens (Balint-Kurti et al. 2010; Manching et al. 2014; McCoy et al. 2019; Singh and Goodwin 2022; Wagner et al. 2020; Wallace et al. 2018). For example, particular phyllospheric bacteria may boost resistance to Cochliobolus*. heterostrophus* infection (Balint-Kurti et al. 2010). That study also correlated high bacterial diversity with SLB susceptibility in the field. Manching et al. (2014) found that severe Southern leaf blight disease is correlated with decreased species richness of epiphytic bacteria. Phyllospheric fungal communities have also been analyzed in fish-eye and marginless tar spot lesions in corn (McCoy et al. 2019).

Despite all these studies, it is not clear whether microbiome structure and disease resistance are directly related or independent processes. Extensive analysis of microbial variation and its correlation to disease, including functional characterization of beneficial microbes, is required to predict and mitigate disease progression. Host disease resistance could be enhanced by the action of antagonist species and/or the net competitiveness of a highly diverse microbiome. Beneficial and/or antagonistic microbes from the corn microbiome may contribute to host resistance by directly inhibiting or suppressing the growth of pathogens or by secreting antimicrobial secondary compounds (reviewed in Morelli et al. 2020; Singh and Goodwin 2022; Trivedi et al. 2020).

As mentioned above, the association of phyllosphere community diversity with disease resistance has been studied previously to some extent (Balint-Kurti et al. 2010; Manching et al. 2014; McCoy et al. 2019; Wagner et al. 2020; Wallace et al. 2018). However, connections between the phyllosphere microbiome and tar spot disease severity have never been investigated. Because the tar spot pathogen exists almost exclusively on the leaf, it and other biotrophic pathogens may be more affected by foliar microbiomes compared to pathogens that live inside host cells. The purpose of this research was to test the hypothesis that microbial community structure differs significantly between resistant and susceptible inbred lines of corn after infection by *P. maydis*. This was accomplished by using Illumina Miseq sequencing to characterize and census the bacteria and fungi associated with resistant and susceptible inbred lines by sequencing the bacterial 16S ribosomal ribonucleic acid (rRNA) gene and the fungal 18S rRNA gene and internal transcribed spacer (ITS) region. We also revealed the significant differential abundance of bacterial and fungal taxa between resistant and susceptible inbred lines. A thorough understanding of microbial diversity and its correlation to disease between resistant and susceptible inbred lines is potentially useful to identify microbes that affect disease progression. Overall, our findings reveal significant differences in community richness and abundance of bacterial and fungal taxa between resistant and susceptible corn inbred lines after infection by *P. maydis*.

## MATERIALS AND METHODS

### Corn lines and growth conditions

In this study, we analyzed the phyllosphere microbiome of sixteen inbred corn lines that differed in severity of tar spot after the onset of symptoms under natural infection in field conditions. Among 16 inbred lines, five were parental lines of the Nested Association Mapping (NAM) population (CML52, CML69, CML103, TX303 and B97) (McMullen et al. 2009; Yu et al. 2008) and the remaining eleven lines (PI685788, PI658790, PI685806, PI685831, PI685836, PI685915, PI685918, PI685919, PI685920, PI685950, and 4401350) were from the Germplasm Enhancement of Maize (GEM) project (Pollak and Salhuana 2001).). Corn inbred lines were planted in three replicates at a research field trial located at the Pinney Purdue Agricultural Center, Wanatah, Indiana. This location was chosen for the experiment because of high tar spot disease pressure detected in prior years. Because of its obligately biotrophic nature, *P. maydis* cannot be cultured for inoculation, hence all field studies relied on natural infection. Plots were planted on June 8, 2019, with approximately 20 seeds per row with two rows for each line in a randomized complete block design (RCBD) with three replications. Experimental plots were 3.0 m (10 ft) wide and 9.1 m (30 ft) long, consisting of two rows for disease evaluation. At all locations, the previous crop was corn with a history of tar spot. Fungicide treatments were not applied during the experimental period to allow for natural infection. After the onset of tar spot symptoms, scoring was conducted at the late dent reproductive growth stage (R5) in early October.

### Sample collection and processing

Leaves from corn lines that ranged from susceptible to tolerant to *Phyllachora maydis* were collected at fourteen weeks after planting. In total, 48 samples were collected from 16 inbred lines including three replicates. For each replicate, five ear leaves were sampled. Sampled leaves were sprayed and wiped with 70% ethanol, and then dried between sterile paper towels. Six leaf discs uniformly spaced from the base to the tip of each leaf were excised using a sterilized cork borer (18 mm), avoiding the midrib and any dry tissue, so that each replication consisted of 30 leaf discs from 5 individual leaves. Excised leaf discs were immediately transferred to sterile microcentrifuge tubes using sterilized forceps and were stored at -80°C until DNA extraction.

### DNA Extraction

DNA was extracted using the Synergy 2.0 Plant DNA Extraction Kit (OPS Diagnostics, Lebanon, NJ, USA) with some modifications. Briefly, leaf discs were ground in liquid nitrogen and homogenized with 500 μl of Plant Homogenization Buffer with a bead beater at its highest speed for 1 minute. Homogenates were centrifuged at 15,000 × g for 5 min at room temperature to pellet debris. Clear supernatant was transferred to a sterilized microcentrifuge tube and RNA was removed by adding 5 μl of RNase A solution, briefly vortexing to mix and incubating at 37°C for 15 min. The extracted DNA was precipitated with isopropanol and transferred to a silica spin column followed by centrifugation at 8,000 × g for 1 min. The DNA samples were washed twice with 250 μl of ice-cold 70% ethanol followed by centrifugation at 8,000 × g for 1 min and dissolved in 50 μl of molecular biology-grade water.

### MetaVx™ library preparation and Illumina MiSeq sequencing

Library preparations and Illumina MiSeq sequencing were conducted at GENEWIZ, Inc. (South Plainfield, NJ, USA). DNA samples were quantified using a Qubit 2.0 Fluorometer (Invitrogen, Carlsbad, CA, USA) and 30-50 ng of DNA were used to generate amplicons using a MetaVx™ Library Preparation kit (GENEWIZ, Inc., South Plainfield, NJ, USA).

#### 16S method for bacteria

For bacterial microbiomes the V3, V4, and V5 hypervariable regions of prokaryotic 16S rDNA were selected for generating amplicons and subsequent taxonomy analysis. GENEWIZ designed a panel of proprietary primers aimed at relatively conserved regions bordering the V3, V4, and V5 hypervariable regions of bacteria and *Archaea* 16S rDNA. The V3 and V4 regions were amplified using forward primers containing the sequence (5’ to 3’) “CCTACGGRRBGCASCAGKVRVGAAT” and reverse primers containing the sequence “GGACTACNVGGGTWTCTAATCC”. The V4 and V5 regions were amplified using forward primers containing the sequence “GTGYCAGCMGCCGCGGTAA” and reverse primers containing the sequence “CTTGTGCGGKCCCCCGYCAATTC”. First-round PCR products were used as templates for second-round amplicon-enrichment PCR. At the same time, adapters were added to the ends of the 16S rDNA amplicons to generate indexed libraries ready for downstream NGS sequencing on an Illumina Miseq machine.

#### 18S method for Eukaryotes

From 50-100 ng of DNA were used to generate amplicons using a panel of primers designed by GENEWIZ (GENEWIZ, Inc., South Plainfield, NJ, USA). Multiple oligonucleotide primers were designed to anneal to the relatively conserved sequences spanning fungal 18S rDNA hypervariable regions V3, V4, V7 and V8, to amplify fragments smaller than 500 bp. The 3’ part of V3 and full V4 regions were amplified using a forward primer containing sequence “GGCAAGTCTGGTGCC” and reverse primer containing sequence “ACGGTATCTRATCRTC”. The entire V7 and V8 regions were amplified using a forward primer containing sequence “CGWTAACGAACGAG” and reverse primer containing sequence “AICCATTCAATCGG”. In addition to the 18S target-specific sequences, the primers also contain adaptor sequences allowing uniform amplification of the library with high complexity ready for downstream NGS sequencing on the Illumina Miseq platform.

#### ITS method for fungi

From 50-100 ng of DNA were used to generate amplicons using a panel of primers designed by GENEWIZ (GENEWIZ, Inc., South Plainfield, NJ, USA). Multiple oligonucleotide primers were designed to anneal to the relatively conserved sequences spanning fungal ITS regions. The ITS1 region was amplified using a forward primer containing sequence “ACCTGCGGARGGAT” and reverse primer containing sequence “GAGATCCRTTGYTRAA”. The ITS2 region was amplified using a forward primer containing sequence “GTGAATCATCGARTC” and reverse primer containing sequence “TCCTCCGCTTATTGAT”. In addition to the ITS target-specific sequences, the primers also contain adaptor sequences allowing uniform amplification of the library with high complexity ready for downstream NGS sequencing on the Illumina Miseq platform.

DNA libraries were validated with an Agilent 2100 Bioanalyzer (Agilent Technologies, Palo Alto, CA, USA), and quantified using a Qubit 2.0 Fluorometer. DNA libraries were multiplexed and loaded on an Illumina MiSeq instrument according to the manufacturer’s instructions (Illumina, San Diego, CA, USA). Sequencing was performed using a 2x300/250 paired-end (PE) configuration; image analysis and base calling were conducted by the MiSeq Control Software (MCS) embedded in the MiSeq instrument.

### Data processing and analysis: *Construction of a database for microbial rDNA sequences*

Fasta-formatted files for the UNITE database of fungal ribosomal sequences (Nilsson et al. 2019) and the Silva database of bacterial, archaeal, and eukaryotic ribosomal sequences (Quast et al. 2013) were downloaded from https://unite.ut.ee and https://www.arb-silva.de, respectively, on 26 April 2021. A third fasta file was composed of the entire *Phyllachora maydis* genome (Telenko et al. 2020) and ribosomal sequences from *Zea mays* and several fungal taxa of interest, including *Monographella nivalis*, *M. stoveri*, *M. lycopodina*, *Coniothyrium* sp., *Paraphaeosphaeria angularis*, *Fusarium sporotrichioides*, and *Neottiosporina paspali*, from NCBI (https://www.ncbi.nlm.nih.gov/genbank/). These three files were concatenated, and custom Perl scripts reformatted the deflines and purged duplicate entries from the combined file. The utility makeblastdb (Camacho et al. 2008) prepared a blastable database, unitesilvaabf, from the combined file. A file (dedupstandardizedsilvahierarchy.txt) of taxonomic names associated with each accession was produced, including genus, family, order, class, and phylum, wherever possible. A series of seven Perl scripts extracted, formatted, and edited the content of this file, which came from the deflines of the fasta files downloaded from https://unite.ut.ee and https://www.arb-silva.de.

#### Quality filtering

Quality-filtered ribosomal reads were obtained from Genewiz, Inc. (South Plainfield, NJ, USA). There were separate forward and reverse fastq files for each inbred. The QIIME data analysis package (Boylen et al. 2019) was used for 16S rDNA and 18S rDNA data analyses. The forward and reverse reads were joined and assigned to samples based on barcode and were truncated by cutting off the barcode and primer sequences. Quality filtering on joined sequences was performed and sequences which did not fulfill the following criteria were discarded: sequence length >200 bp, no ambiguous bases, mean quality score >= 20. Then the sequences were compared with the reference database (RDP Gold database) using the UCHIME algorithm (Edgar 2016) to detect chimeric sequences, which were then removed.

Quality was confirmed by subjecting 117 files of reads to the HTStream pipeline (https://s4hts.github.io/HTStream/), minus its steps to remove ribosomal sequences and eliminate PCR duplicates. This aggressive trimming pipeline shortened 117 of the reads files by less than 0.3%. Therefore, the reads were used without further trimming or deduplication.

#### Taxonomic annotation and relative abundance analysis

Filtered reads were reformatted to fasta and aligned with blastn (Camacho et al. 2008) against the unitesilvaabf database with an expected value of 1e^-40^ and a one-line output format per hit. The default blastn limits on number of returned reads applied. Therefore, there were frequent ties among two or more taxa for closest hit.

Script countfromunitesilvaabf2.pl was written to count closest hits. It utilized the query sequence file, its blastn output, and the file dedupstandardizedsilvahierarchy.txt of associated taxonomic names. It credited taxa with all hits to their accessions in the unitesilvaabf database. In case of a tie, the hit was divided evenly among all tied taxa, and thus fractional counts were frequent among the total counts for each taxon. Scripts mergecountsbyrank0325.pl and mergecountsbyrank0329.pl each collected counts by taxon at the five levels of phylum, class, order, family, and genus. These two scripts differed in calculation of the output. The former transformed counts to percentage of all the blast hits and transformed percentage to natural log of (1 + percentage) for the subsequent construction of heatmaps. The latter output was used directly for the subsequent construction of barplots and application of DESeq2 to assess significance of population differences. With both scripts, there were separate outputs with and without counts to *Phyllachora maydis*, *Phyllachoraceae*, *Phyllachorales*, or their contribution to counts of *Sordariomycetes* and *Ascomycota*. In addition to a combined output for all taxa, both scripts also output 10 tables of the 30 most-abundant phyla, classes, orders, families, and genera within prokaryotes and eukaryotes. Eukaryotic heatmaps and barplots were prepared from these tables both with and without *Phyllachora* counts; the latter were used to investigate the relative effect of *Phyllachora* on populations of the other taxa. Heatmaps were produced with the R functions “heatmap.2” in package “gplots” version 3.1.1 (Warnes et al. 2020) and “viridis” in package “viridis” version 0.6.2 (Garnier et al. 2021). Barplots were constructed using function geom_rect in R package “ggplot2” version 3.3.3 (Wickham 2016), with boundaries calculated with Perl script buildpctbarplot0330.pl. Because the rectangle areas were directly proportional to the fraction of counts among the top 30 but not all of the taxa, the ranking of inbreds for fraction of *Phyllachora* reads differed from that used for heatmaps.

#### Population diversity analysis (alpha and beta diversity)

Population statistics were calculated separately for genera, families, orders, classes, and phyla, and for prokaryotes, eukaryotes with *Phyllachora*, and eukaryotes without *Phyllachora*. Six indices of taxonomic alpha diversity (Chao1, ACE, Shannon, Simpson, inverse Simpson, and Fisher) were calculated with integerized genus-level counts using function “estimate_richness” in R package “phyloseq” version 1.22.3 (McMurdie and Holmes, 2013). The Sorensen and Bray-Curtis indices of beta diversity were calculated with function “vegdist” in R package “vegan” version 2.5-7 (Oksanen et al. 2020). The 18 sets of alpha diversity index values (six indices, each replicated thrice) were subjected to one-way ANOVA with R function “aov”, and pairwise estimates of significance were calculated with Tukey’s Honest Significant Difference test (R function “TukeyHSD”). The significance of differences was graphed with R package “ggplot2” version 3.3.3 using two Perl scripts that used (1 – p), where p is the TukeyHSD *p*-value of any of the alpha diversity indices for any pair of inbreds. The Perl scripts found a minimum-spanning tree of (1 – p) over all combinations of inbreds and set up a circular display where cross links denoted greater similarity in the specified alpha-diversity index.

Statistical significance of population variation was also tested with R package “DESeq2” version 1.30.1 (Love et al. 2014). Counts were rounded to the nearest integer. The test compared the five inbreds with the lowest frequency of *Phyllachora* (CML103, CML69, CML52, PI685831, PI685915) to the five inbreds with the highest frequency of *Phyllachora* (PI685788, TX303, PI685790, 4401350, PI685920) reads. The remaining six inbreds were excluded from this analysis. Pearson correlation coefficients between the relative frequencies of 20 most-abundant bacterial and fungal taxa with increasing *P. maydis* reads were calculated by using the R function cor.test. Furthermore, the Pearson correlation column of Table 3 was derived by calculating these quantities for each combination of inbred and replicate: A, the count of *P. maydis* reads; Bi, the count of reads for each top-20, non-Phyllachora maydis fungal taxon i; D, the total count of fungal reads; Ci, the read count of each non-HG917238 (*Acinetobacter sp*.), top-20 bacterial taxon i; and E, the read count of all bacterial taxa except HG917238 (*Acinetobacter sp*.), which was treated as maize chloroplast contamination because of the abundant secondary hits at 88-91% identity to cyanobacteria and chloroplasts and the dearth of additional hits to Acinetobacter genomes. Then Bi / (D – A) was calculated for each fungal taxon i, and Ci / E was calculated for each bacterial taxon i. The ratios of Bi / (D – A) and Ci / E were compared to the ratio A/D using the R function cor.test. *p*-values were adjusted for false discovery rate by the method of Benjamini and Hochberg in R function *p*.adjust. The objective was to find any significant change in the top 20 taxa that could indicate a response other than simple, physical displacement of other taxa by *P. maydis*.

## RESULTS

### Inbred corn lines that varied for tar spot severity under natural infection were used for Miseq sequencing

Disease scoring data showed that inbreds TX303, B97, PI685788, PI658790, PI685831, PI685836, PI685915, PI685918, PI685920, PI685950, and X4401350 were susceptible whereas CML103, CML69, CML52, PI685806 and PI685919 were resistant towards *P. maydis* (Table 1). However, the comparison of Miseq sequencing revealed that inbred lines PI685806, PI685919 and CML52 had more than 20% *P. maydis* reads even though they looked resistant based on disease scores (Table 1). Hence, based on both visual analysis and Miseq read counts, we considered only CML103 and CML69 as resistant lines and the other fourteen as susceptible.

**Table 1.**
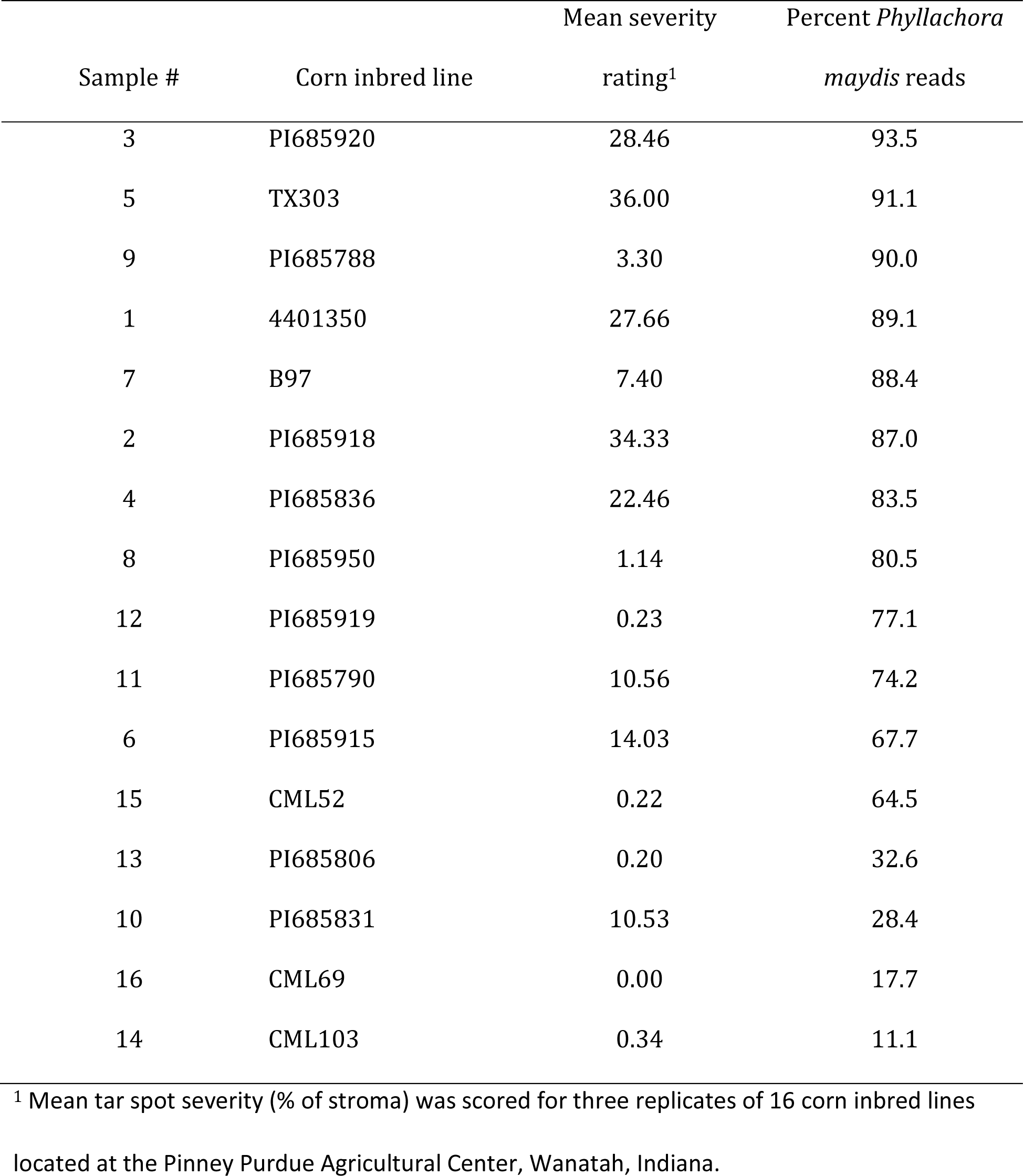
Disease scores and percent *Phyllachora maydis* sequence reads for sixteen inbred lines of corn (*Zea mays* ssp. *maydis*) chosen from the parents of the Nested Association Mapping (NAM) and Germplasm Enhancement of Maize (GEM) genetic populations.

| Sample # | Corn inbred line | Mean severity | Percent *Phyllachora* |
| --- | --- | --- | --- |
|  |  | rating<sup>1</sup> | *maydis* reads |
| 3 | PI685920 | 28.46 | 93.5 |
| 5 | TX303 | 36.00 | 91.1 |
| 9 | PI685788 | 3.30 | 90.0 |
| 1 | 4401350 | 27.66 | 89.1 |
| 7 | B97 | 7.40 | 88.4 |
| 2 | PI685918 | 34.33 | 87.0 |
| 4 | PI685836 | 22.46 | 83.5 |
| 8 | PI685950 | 1.14 | 80.5 |
| 12 | PI685919 | 0.23 | 77.1 |
| 11 | PI685790 | 10.56 | 74.2 |
| 6 | PI685915 | 14.03 | 67.7 |
| 15 | CML52 | 0.22 | 64.5 |
| 13 | PI685806 | 0.20 | 32.6 |
| 10 | PI685831 | 10.53 | 28.4 |
| 16 | CML69 | 0.00 | 17.7 |
| 14 | CML103 | 0.34 | 11.1 |

<sup>1</sup> Mean tar spot severity (% of stroma) was scored for three replicates of 16 corn inbred lines
located at the Pinney Purdue Agricultural Center, Wanatah, Indiana.

### Structure and diversity of the bacterial and fungal communities in resistant and susceptible lines

Taxonomic profiling was performed by mapping bacterial and fungal reads to the reference databases (SILVA 16S v.8.3 for bacterial reads and UNITE fungal ITS v138.1 for fungal reads) to analyze the composition and diversity of microbial communities in tar spot-resistant and -susceptible samples. The Illumina Miseq sequencing of 48 samples including 3 replicates resulted in 10,868,490 bacterial reads and 16,217,432 fungal reads, with on average 113,213 bacterial reads per sample and 172,525 fungal reads per sample. In total, the reads could be characterized into 748 bacterial and 1693 fungal genera across all corn lines (Table 2). The dominant bacterial families were *Moraxellaceae* and *Beijerinckiaceae* in all inbred lines. Additional families such as *Erwiniaceae*, *Sphingomonadaceae*, *Kineosporiaceae*, *Geodermatophilaceae*, *Spirosomaceae*, *Hymenobacteraceae*, *Nocardioidaceae*, *Rhizobiaceae* and *Enterobacteriaceae* were also detected to some extent in all inbred lines (Fig. 1A). The dominant bacterial genera were *Moraxellaceae gen., Erwiniaceae gen., Methylobacterium*, *Beijerinckiaceae gen., Pantoea, Methylorubrum* and *Sphingomonas* whereas *Quadrisphaera*, *Nocardioides*, *Klenkia* and *Spirosoma* were also detected (Fig. 1B). Interestingly, families *Erwiniaceae, Sphingomonadaceae, Rhizobiaceae,* and *Microbacteriaceae* and genera *Methylobacterium* and *Pantoea* were highly abundant on most of the susceptible lines. Additionally, family *Pseudomonadaceae* and genus *Pseudomonas* were highly abundant on PI685831 only. At the phylum level, *Proteobacteria*, *Actinobacteriota*, and *Bacteroidota* were the dominant bacterial phyla on all lines (Supplementary Fig. S1).

**Fig. 1.**
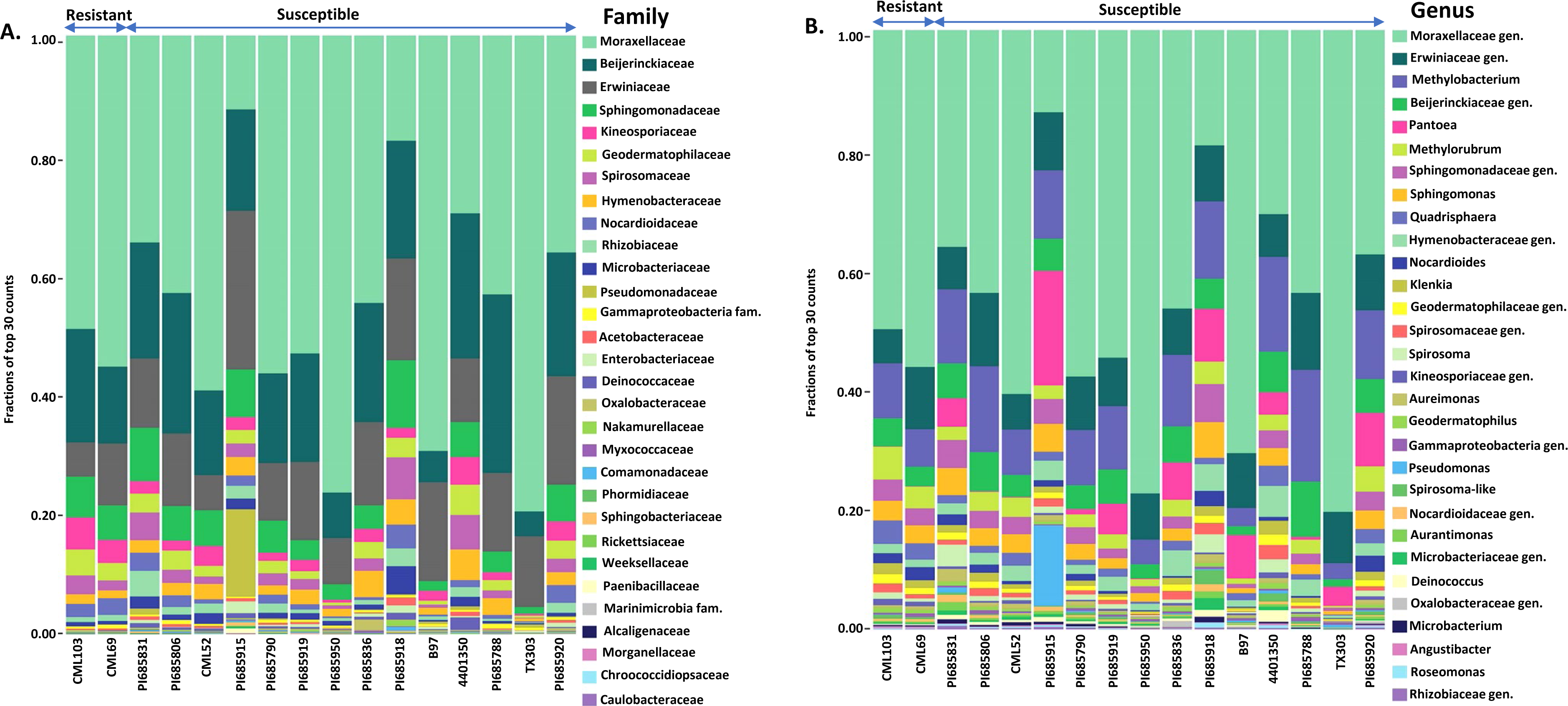
The 30 most abundant bacterial families (**A**) and genera (**B**) in 16 inbred lines of corn sorted from left to right as resistant or susceptible based on the number of *Phyllachora maydis* sequence reads. The values represent relative abundances averaged over three replications, and each color specifies a listed taxon.

**Table 2.** Numbers of prokaryotic and eukaryotic taxa that appeared in at least one sample of microbiome sequences from 16 inbred lines of corn that differed in resistance to tar spot caused by the fungal pathogen *Phyllachora maydis*, by taxonomic level

| Taxonomic level | Prokaryotic taxa | Eukaryotic taxa |
| --- | --- | --- |
| Phylum | 36 | 33 |
| Class | 62 | 74 |
| Order | 151 | 156 |
| Family | 253 | 357 |
| Genus | 748 | 1693 |

The most abundant fungal families were *Phyllachoraceae*, *Cladosporiaceae*, *Didymellaceae*, and *Pleosporaceae*, while *Coniothyriaceae*, *Bulleribasidiaceae*, *Phaeosphaeriaceae*, *Spiculogloeaceae* and unidentified family in order *Entylomatales* were also detected in minor numbers (Fig. 2A). The most abundant genera were *Phyllachora*, *Cladosporium*, those in the order *Pleosporales*, *Coniothyrium*, *Bipolaris* and *Ascochyta*, with *Phyllozyma* and *Tilletiopsis* being detected at low levels (Fig. 2B). Interestingly, family *Phyllachoraceae* and genus *Phyllachora* were detected in all inbreds. Numbers of *Phyllachora* reads mostly corresponded to the disease score exhibited by the inbred lines, with PI685920, PI685918, 4401350, PI685836, TX303, PI685915, B97, PI685950, PI685788, PI685831 and PI685790 having more than 20% *Phyllachora* reads and CML103 and CML69 having the reads lower than 20% (Fig. 2A, 2B and Table 1). In contrast, *Phyllachora* read percents in inbred lines PI685919, PI685806 and CML52 were not consistent with the visual disease assessment (Table 1). Furthermore, family *Cladosporiaceae* in the *Ascomycetes*, order Entylomatales in the class *Exobasidiomycetes* (*Basidiomycota*), family *Trichocomaceae* in the class *Eurotiomycetes*, families *Tremellaceae* and *Rhynchogastremaceae* in the class *Tremellomycetes* were dominant on resistant lines CML103 and CML69 (Fig. 2A). The families *Pucciniaceae* and *Leptosphaeriaceae* were most abundant on the inbred lines PI685915, B97 and TX303, whereas family *Didymosphaeriaceae* was most abundant on susceptible inbred lines PI685836, PI685918, 4401350 and PI685788 (Fig. 2A). Intrestingly, family *Dothideaceae* and genus *Neottiosporina* were predominantly detected only on PI685836 and PI685918. At the genus level, *Cladosporium* and *Tilletiopsis* predominated on resistant lines CML103 and CML69 (Fig. 2B). *Bipolaris* was the most-abundant genus on the lines PI685831, PI685806 and PI685915, whereas *Puccinia* and *Sphaerellopsis* (a mycoparasite of *Puccinia* species) were most abundant on inbreds PI685915, B97 and TX303 (Fig. 2B). In addition to the unidentified fungi phylum, the fungal phyla *Ascomycota* and *Basidiomycota* were the dominant fungi on all inbred lines (Supplementary Fig. S2). These findings indicate that the phyllosphere microbiome of the sixteen inbred corn lines evaluated had low diversity in terms of bacterial and fungal distribution, with only a few dominant taxa identified. Although sample-to-sample variability is still high, the distribution of dominant bacterial and fungal taxa in all inbreds followed the same basic trend.

**Fig. 2.**
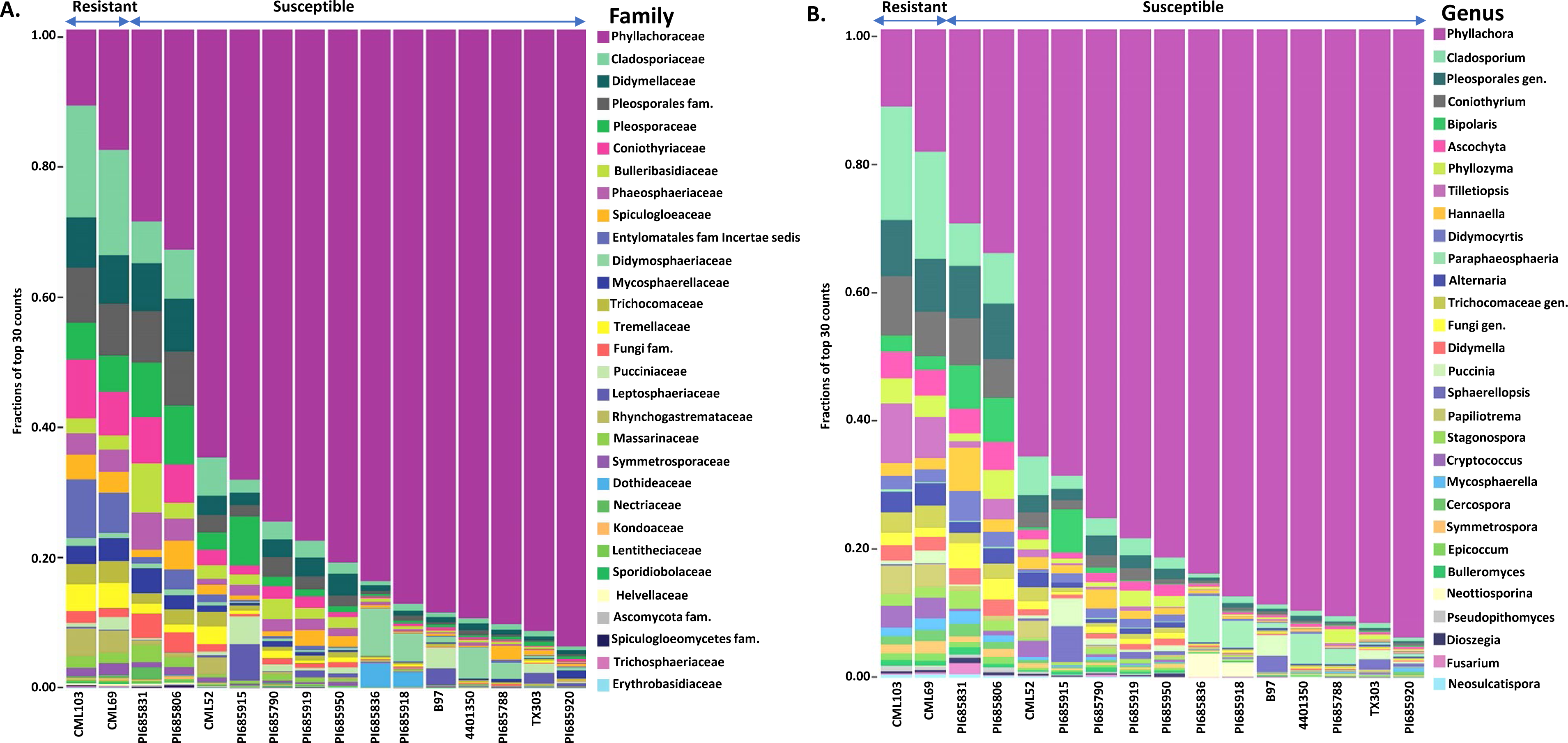
The 30 most abundant fungal families (**A**) and genera (**B**) in 16 inbred lines of corn sorted from left to right as resistant or susceptible based on the number of *Phyllachora maydis* sequence reads. The bars represent the relative abundances averaged over three replications, and each color specifies a listed taxon.

### Alpha (*α*) and beta (β) diversities showed differences in bacterial and fungal communities between resistant and susceptible corn lines

The *α* diversity is a statistic that is used to estimate richness (number of taxa), evenness (distribution of frequencies of the taxa), or both within a sample. For the *α* diversity analysis, four metrics were used: the Chao1 index (richness), ACE (richness), Shannon index (richness and evenness), and the Simpson index (richness and evenness). Higher values of the Chao1 or ACE indices represent higher species richness of the microbiota within a community (Chao 1984; Chao et al. 1993), while higher values of the Shannon index represent samples with more taxa with similar frequencies among the microbiota within a community. In contrast, lower values of the inverse Simpson index represent higher diversity (richness and evenness as with the Shannon statistic) of the microbiota within a community (Simpson 1949). When compared together, resistant and susceptible corn lines showed some variation in α diversity (richness) for both bacterial and fungal communities (Fig. 3). The bacterial community richness (ACE: p-value ≤ 0.05; and Chao: p-value ≤ 0.05) of the two resistant lines CML103 and CML69 was higher than those of fourteen susceptible inbred lines (Fig. 3, A and B). Furthermore, the fungal community richness (ACE: p-value ≤ 0.05; and Chao: p-value ≤ 0.05) also was higher in the two resistant lines CML103 and CML69 compared to the susceptible inbreds, except for PI685831 and PI685806, which had higher fungal community richness (Fig. 3, C and D). Replications were generally fairly consistent, especially for samples with low taxon richness (Fig. 3). The most striking variation occurred for inbred PI685806, where two replications showed the highest fungal richness among all samples but the third was lower (Fig. 3, C and D).

**Fig. 3.**
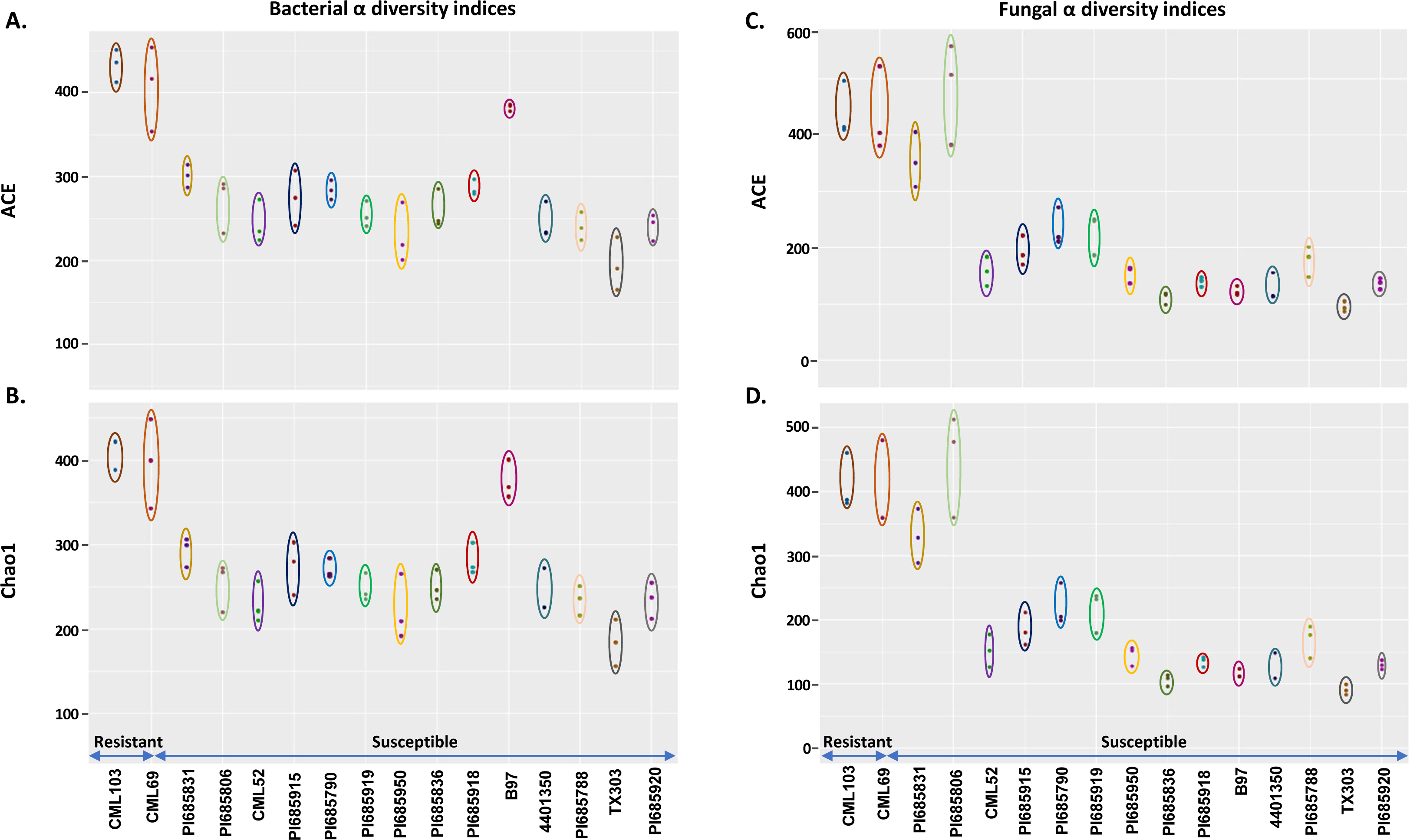
Bacterial and fungal alpha diversity (richness) measures for the ACE and Chao indices for bacteria (**A** and **B**) and fungi (**C** and **D**) observed on corn inbreds that are resistant or susceptible to *Phyllachora maydis* based on increasing proportion of pathogen reads in the samples. The ACE and Chao indices reflect the microbiota abundance (richness) in the three replications per inbred. The colored ovals coordinate the values of the same inbred in different panels. Significant differences were calculated by one-way ANOVA with R function “aov”, and pairwise estimates of significance were calculated with Tukey’s Honest Significant Difference test (R function “TukeyHSD”) (*p* ≤ 0.05) (Table S1).

When evenness was included in the calculations there was no clear difference in diversity of bacterial communities among the tolerant and susceptible lines as shown by the Shannon and Simpson indices (Fig. 4). Except for the susceptible inbred lines PI685790 and TX303, which were significantly lower, the remaining fourteen lines showed similar levels of bacterial community diversity (Fig. 4, A and B). However, there was a clear difference in diversity of fungal communities among the resistant and twelve of the susceptible lines (CML52, PI685915, PI685790, PI685919, PI685950, PI685836, PI685918, B97, 4401350, PI685788, TX303, and PI685920) which had lower fungal community diversities than those of resistant lines CML103 and CML69 or the other less susceptible lines (Fig. 4, C and D). Based on these data we can conclude that resistant lines CML103 and CML69 contained higher levels of species richness in their bacterial and fungal microbial communities compared with susceptible lines, but no significant differences in evenness of bacterial communities were observed between the resistant and susceptible lines. For the fungal microbiota, the two most resistant lines based on *Phyllachora* reads had higher evenness compared to the twelve most highly susceptible lines (Fig. 4, C and D). The replications were mostly very consistent with a few exceptions. For example, inbred CML69 had the highest and very uniform evenness for fungal taxa (Fig. 4, C and D), but had the widest variation among replications for all lines when richness was included for both bacterial and fungal diversity (Fig. 3, A, B, C and D), indicating a more uneven frequency distribution among replications of the reduced number of species.

**Fig. 4.**
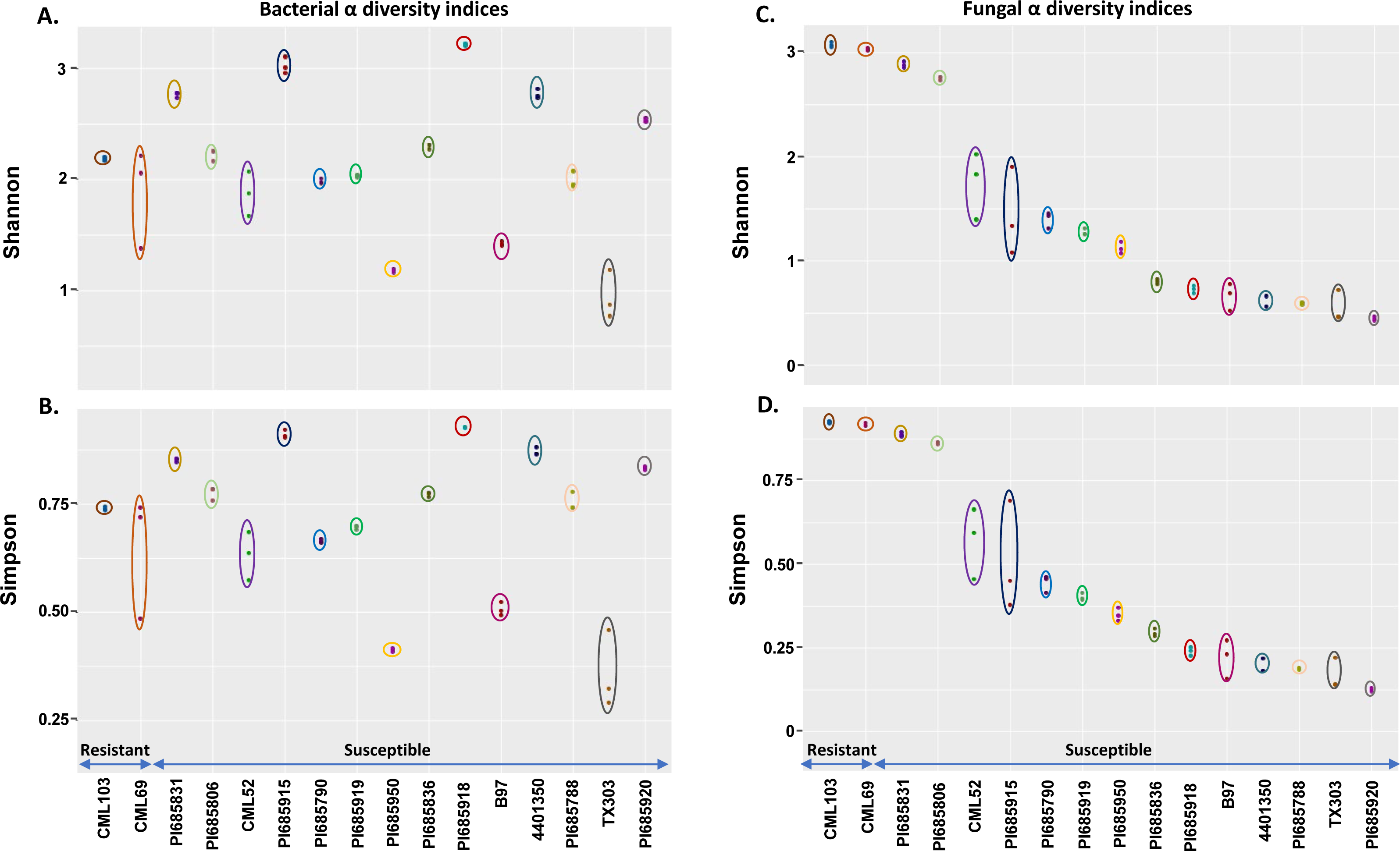
Bacterial and fungal alpha diversity when considering both richness and evenness as measured by the Shannon and (B) Simpson indices for bacteria (**A** and **B**) and fungi (**C** and **D**) observed on corn inbreds that are resistant or susceptible to *Phyllachora maydis* based on increasing proportion of pathogen reads in the samples. The Shannon and Simpson indices reflect the diversity of microbial taxa in the three replications per inbred. The colored ovals coordinate the values of the same inbred in different panels. Significant differences were calculated by one-way ANOVA with R function “aov”, and pairwise estimates of significance were calculated with Tukey’s Honest Significant Difference test (R function “TukeyHSD”) (*p* ≤ 0.05) (Table S1).

Estimates of beta diversity enable comparisons among microbial communities, considering both the number of distinct microorganisms in the sample and their phylogenetic relatedness, i.e., differences in their specific species composition. The beta diversities of the microbial communities between the tolerant and susceptible lines were compared using the Sorensen’s beta (Sorensen 1948) and Bray-Curtis dissimilarity (Bray and Curtis 1957) metrics. The Sorensen’s beta is based on presence or absence of taxa, whereas Bray-Curtis dissimilarity considers abundances of taxa in two samples. Both measures range between 0 and 1, with 0 indicating no similarity and 1 indicating the samples contain the same species at the same frequencies. Analysis of the beta diversity using Sorensen’s metric (0.170 for fungi and 0.111 for bacteria; ANOVA p-value < 2e^-16^ for both) showed the presence of a similar suite of species in all inbred lines. Furthermore, the Bray-Curtis dissimilarity metric (0.571 for fungi, and 0.464 for bacteria; ANOVA p-value<2e-16 for both) showed that the microbial populations on the susceptible lines had differentially abundant species with increasing *Phyllachora* populations compared to the tolerant lines with fewer *Phyllachora* read counts.

### Identification of differential taxon abundance between resistant and susceptible lines

To identify the bacterial and fungal taxa associated with resistant and susceptible lines, we performed differential abundance analysis using DESeq2 as 0.05 false discovery rate as the threshold of significance. At the family level, *Geodermatophilaceae*, *Kineosporiaceae* and *Nocardioidaceae* and on a genus level, *Nocardioides, Geodermatophilus*, *Kineosporiaceae* genus*, Angustibacter*, *Methylorubrum*, and *Quadrisphaera* were the most abundant prokaryotic taxa in the resistant inbred lines (Fig. 5A). In contrast, the prokaryotic genera *Spirosoma*, *Aureimonas*, *Deinococcus*, *Pseudomonas,* and *Pantoea*, were the most abundant ones in the susceptible lines (Fig. 5A). Interestingly, *Klenkia*, *Sphingomonas and Methylobacterium* were abundant in both resistant and susceptible lines.

**Fig. 5.**
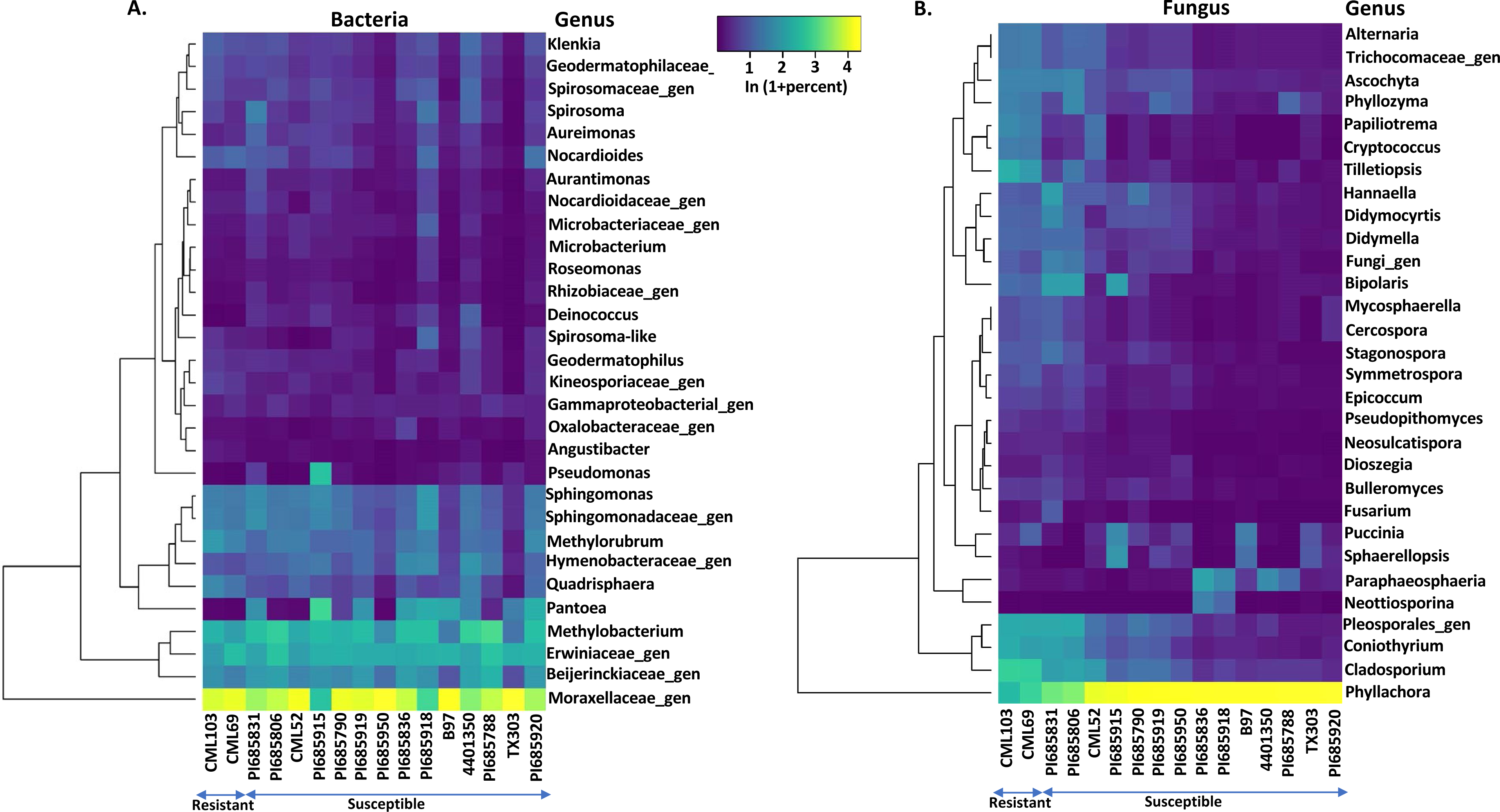
Heat map showing the relative abundances of bacterial and fungal genera in the corn leaf microbiome. Relative abundances of the 30 most abundant **(A)** bacterial and **(B)** fungal genera at p value ≤ 0.05 in 16 inbred corn lines sorted from left to right as resistant or susceptible based on the number of *Phyllachora maydis* sequence reads.

Fungal families *Cladosporiaceae*, *Pleosporaceae*, *Didymellaceae*, *Trichocomaceae*, *Tremellaceae*, *Rhynchogastremataceae* and a family in the *Entylomatales* were more abundant in resistant lines whereas families *Phyllachoraceae*, *Pucciniaceae*, *Dothideaceae* and *Leptosphaeriaceae* were more abundant in susceptible lines. Fungal genera *Alternaria, Trichocomaceae* genus*, Ascochyta, Phyllozyma, Papiliotrema*, *Cryptococcus*, *Tilletiopsis*, *Bipolaris*, *Stagonospora*, *Symmetrospora, Epicoccum, Bulleromyces, Pleosporales* genus*, Coniothyrium* and *Cladosporium* were the most abundant in the resistant lines (Fig. 5B). The genera *Phyllachora*, *Sphaerellopsis*, *Puccinia* and *Didymella* were highly abundant in susceptible lines, suggesting these taxa were possibly associated with tar spot (Fig. 5B).

### Tar spot disease severity level altered the phyllosphere microbiome

*Phyllachora maydis* reads were observed in all inbred lines irrespective of their severity level. To check whether *P. maydis* status influenced the phyllosphere microbiome, the relative abundance of taxa in resistant and susceptible inbreds was investigated with DESeq2 (Love et al. 2014), which generated lists of taxa, mean counts, log 2-fold-changes, and *p*-values in the same way as it treats differential gene expression. Positive log-2-fold change indicated an increase in susceptible versus tolerant inbreds. The following taxa were noted as significantly responding to increasing *Phyllachora* at *p* ≤ 0.05, and the taxa are listed in ascending order of *p*-value, thus from most to least significantly responding within the limit of *p*≤0.05. Bacterial family *Rhodanobacteraceae* and genera *Methylobacterium*, *Luteibacter*, *Kineosporia*, genera in the family *Methylobacteriaceae*, *Fibrella*, *Loriellopsis*, *Kluyvera*, *Klugiella*, *Novosphingobium* and *Massilia* increased in the susceptible inbred lines. Bacterial genera *Pseudokineococcus*, unidentified genera in the family *Aurantimonadaceae*, *Fulvimarina* and *Ornithinimicrobium* decreased in the susceptible inbreds.

The eukaryotic phylum *Ascomycota* was more abundant in susceptible lines whereas phyla *Basidiomycota* and *Mucoromycota* were less abundant. Fungal families *Phyllachoraceae*, *Trimorphomycetaceae*, *Bulleraceae*, *Spiculogloeaceae*, *Corticiaceae* and *Sporormiaceae* were more abundant in susceptible inbreds, while *Helvellaceae* and *Cladosporiaceae* decreased. The genera *Phyllachora*, *Saitozyma*, *Epicoccum*, *Genolevuria*, *Phyllozyma*, *Ascochyta*, *Limonomyces*, *Paraphaeosphaeria*, *Preussia*, *Cercospora*, *Scheffersomyces*, *Holtermanniella* and *Tetraplosphaeria* increased in susceptible lines and possibly were associated with tar spot. Meanwhile, *Leptospora*, *Paraophiobolus*, *Cladosporium*, *Pseudozyma*, *Bipolaris*, and *Ustilago* decreased in the susceptible lines. Overall, DESeq2 analysis revealed that *Phyllachora* status had a significant impact on abundance resulting in either a reduction or increase of bacterial and/or fungal microbial taxa. It seems likely that *P. maydis* was associated with the bacterial and fungal taxa that were present in enhanced relative abundance in susceptible inbreds in response to increased *Phyllachora* reads.

To highlight potential links between the bacterial and fungal taxa with increasing *P. maydis* reads, Pearson correlation coefficients were calculated between the relative abundances of the 20 most abundant bacterial (Table 3) and fungal (Table 4) taxa. Intriguingly, correlation analysis revealed more negative correlations than positive ones, at least for the most abundant taxa in the bacterial and fungal communities, suggesting that the taxon is mutually exclusive. Among the 20 most abundant bacterial taxa, only four bacterial species, *Pantoea ananatis*, *Methylobacterium radiotolerans*, *Methylobacterium* komagatae, and *Methylobacterium organophilum*, were statistically and positively correlated with *P. maydis* reads, while nine bacterial species including *Sphingomonas melonis*, *Geodermatophilus* sp., *Sphingomonas hankookensis*, *Klenkia brasiliensis*, *Methylorubrum extorquens*, *Nocardioides* lentus, *Quadrisphaera* setariae, *Quadrisphaera granulorum*, and *Spirosoma oryzae*, were significantly negatively correlated with *P. maydis* reads (Table 3). In addition, for fungal species, only *Paraphaeosphaeria michotii* was positively correlated with *P. maydis* reads, whereas *Papiliotrema flavescens*, *Alternaria* sp., *Amphinema* sp., *Neosetophoma xingrensis*, *Neottiosporina paspali*, *Coniothyrium* sp., *Cercospora* beticola, *Symmetrospora symmetrica*, *Cercospora zeae-maydis*, *Tilletiopsis washingtonesis*, *Didymella coffeaearabicae*, *Stagonospora* sp., *Phyllozyma linderae*, *Bullera alba*, *Dioszegia* zsoltii and *Bipolaris eleusines* were statistically and negatively correlated with *P. maydis* reads (Table 4). The overall findings showing more negative correlations with *P. maydis* reads indicates a high possibility for increased ecologic niche differentiation because of a high proportion of competition among bacterial and fungal communities.

**Table 3.**
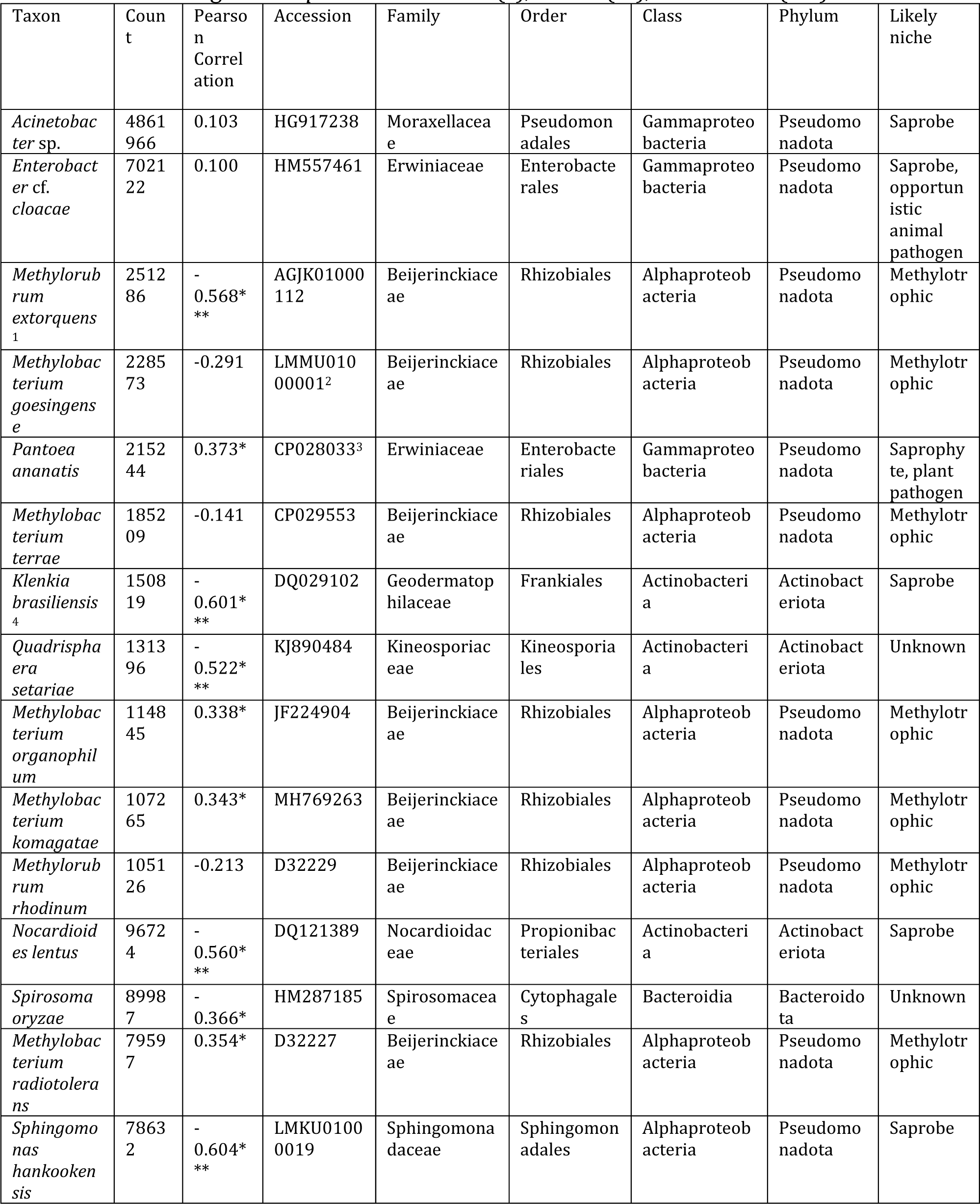
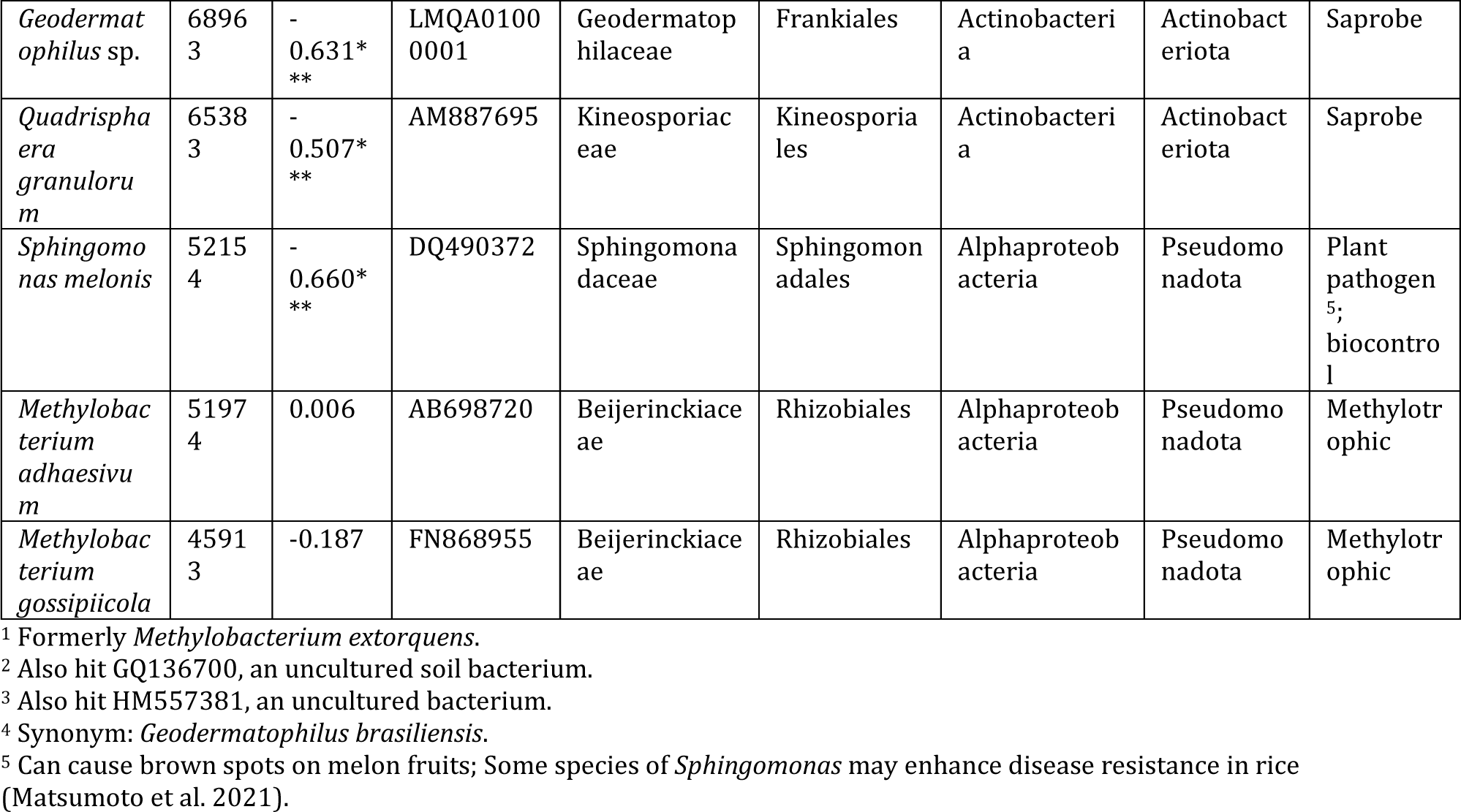
The 20 most common bacterial taxa over all inbred lines and replications identified to most likely species, and their correlations with the percent of *Phyllachora maydis* reads. Negative correlations imply mutual exclusivity of the taxon and *P. maydis*. Asterisks indicate significant *p*-values at ≤ 0.05 (*), ≤ 0.01 (**), and ≤ 0.001(***).

**Table 4.**
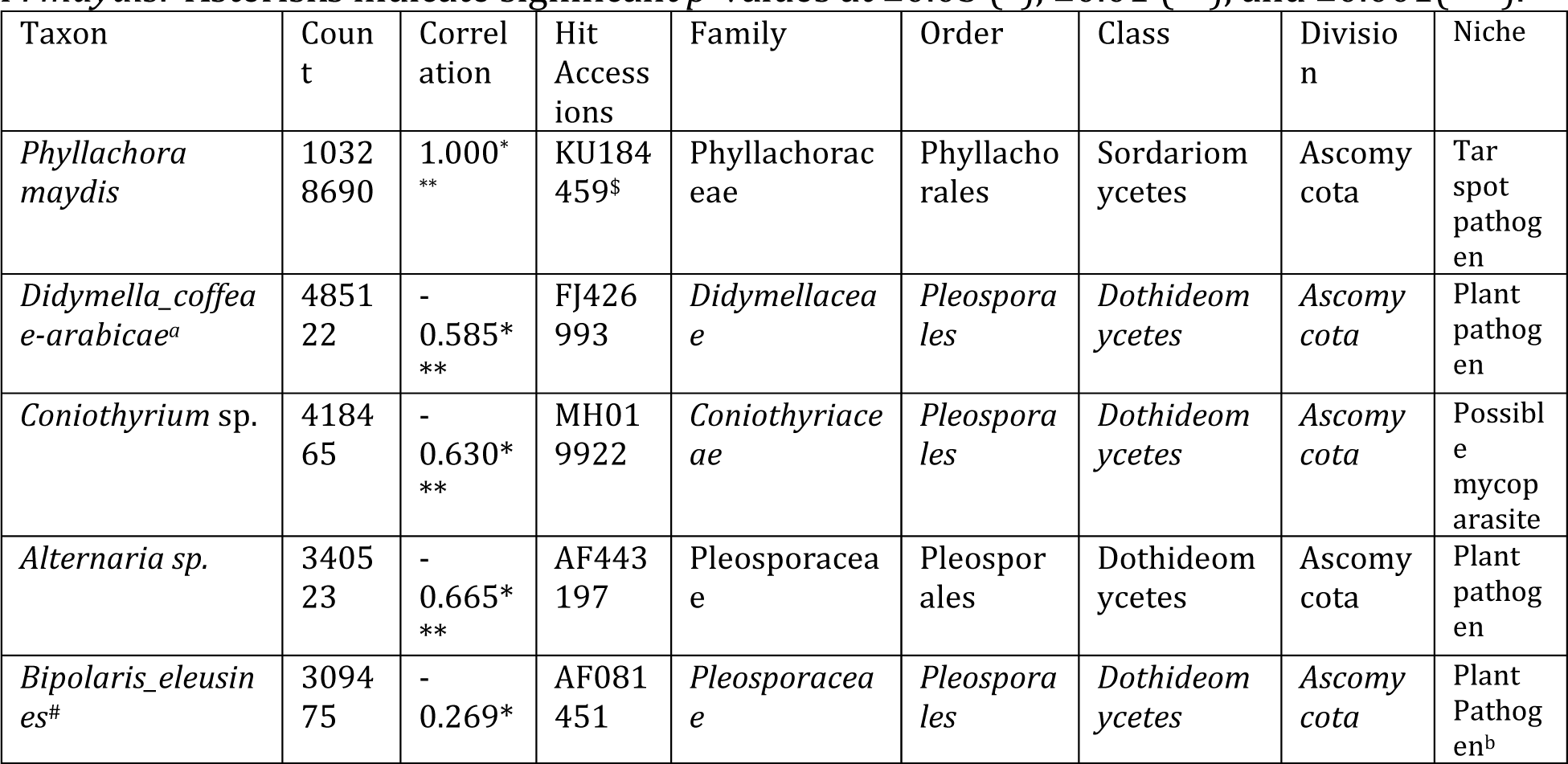
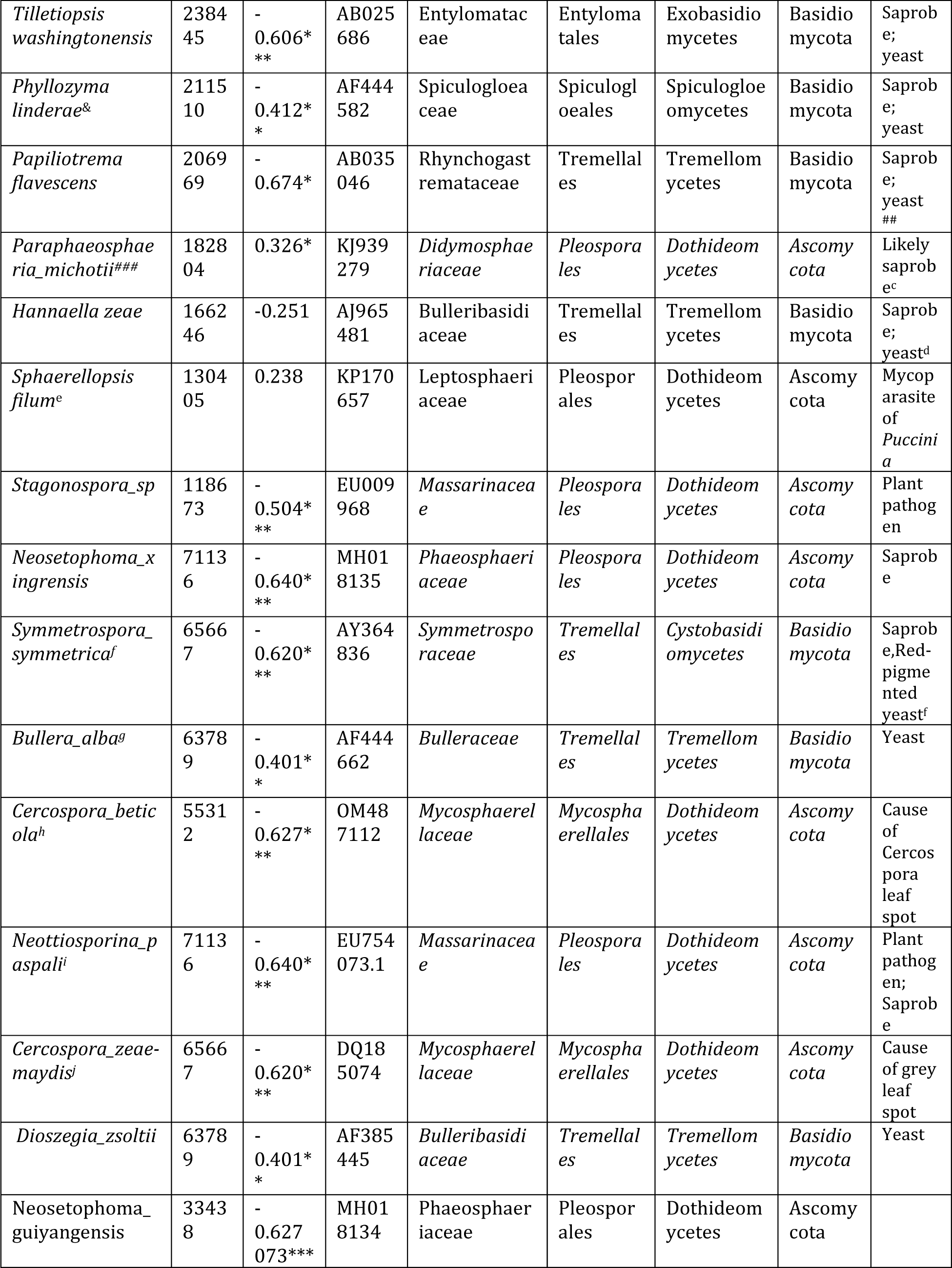
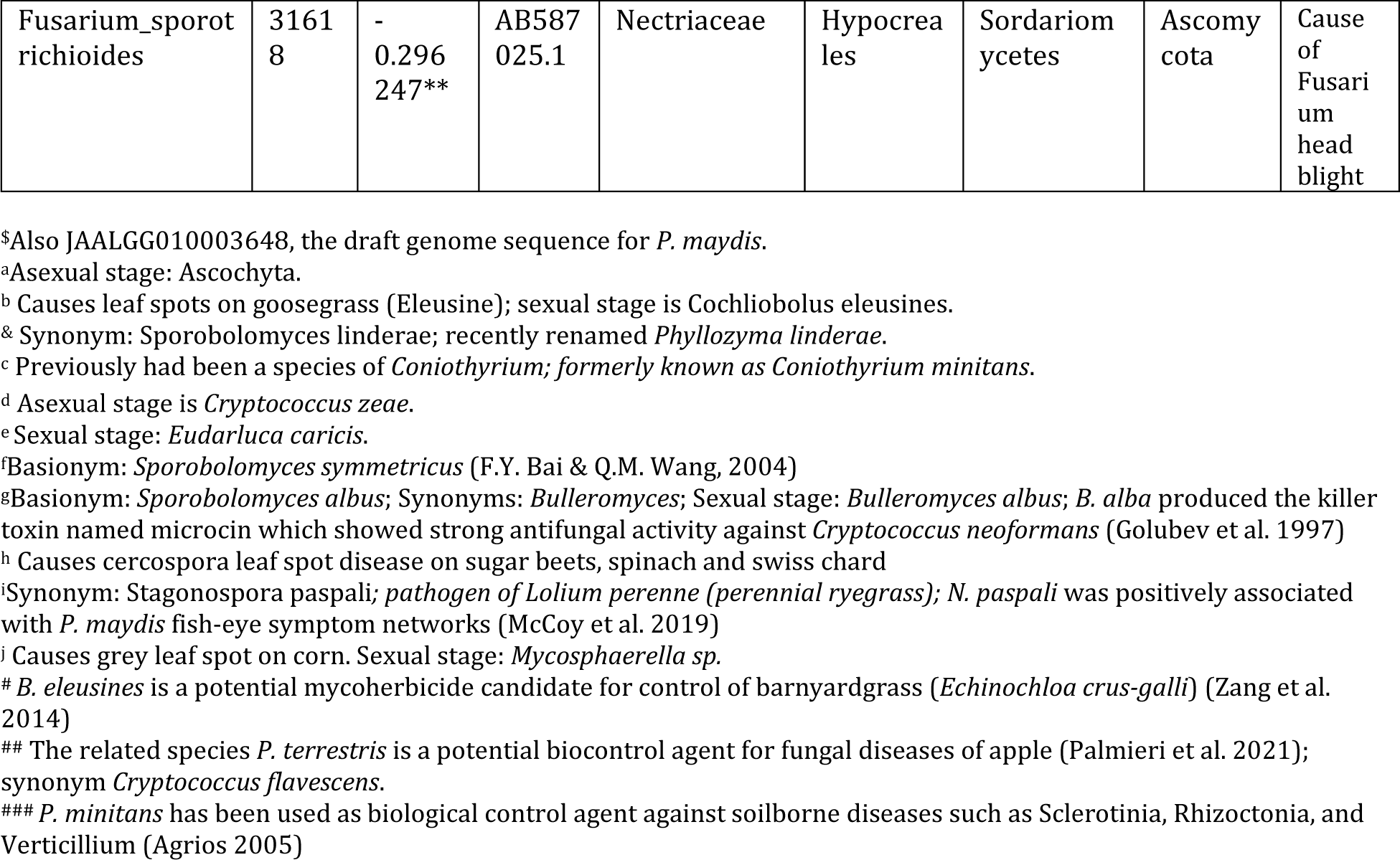
The 20 most abundant fungal species encountered over all inbreds. Count is the sum of read counts over all inbreds and replicates. Pearson’s correlation coefficient r was calculated from abundance of each taxon versus abundance of *Phyllachora maydis* in the same inbred and replicate. Negative correlations imply mutual exclusivity of the taxon and *P. maydis*. Asterisks indicate significant *p*-values at ≤0.05 (*), ≤0.01 (**), and ≤0.001(***).

In addition to tar spot, the sequence data also indicated corn lines that were susceptible to other common fungal pathogens. For example, inbreds PI685831, PI685915 and PI685806 were susceptible to southern corn leaf blight caused by *Bipolaris maydis*, line PI685920 to gray leaf spot (*Cercospora spp*), and lines PI685915, B97 and TX303 to rust (*Puccinia* spp.) (Fig. 5B). Interestingly, the inbreds with high read numbers from rust pathogens also had high sequence reads for *Sphaerellopsis* sp., a known mycoparasite of *Puccinia* spp. Therefore, the microbiome sequences provided a good overall picture of the fungi interacting with each other and their host substrate in the corn phyllosphere.

## DISCUSSION

The phyllosphere can contain a wide variety of beneficial, pathogenic, and commensal microbes (Lindow and Brandl 2003). Although phyllospheric microbial communities in corn have been studied to some extent (Singh and Goodwin 2022), comparatively little is known about the structure, diversity, and community dynamics of this microbiome or its interactions with the host (Penuelas and Terradas 2014; Singh and Goodwin 2022). Further analysis of the foliar microbiome is critical for understanding the types of pathogens present and evaluating the potential of other microbes to predict or prevent disease progression.

To test the hypothesis that microbiome composition varied in response to *P. maydis* infection in resistant and susceptible lines, we sequenced rRNA amplicons to investigate bacterial and fungal populations in 16 corn inbreds that varied in tar spot severity. We found that fungal taxa and populations responded more than bacteria to increasing percentage of *P. maydis* sequence counts. Alpha-diversity indices for bacterial and fungal microbiomes differed between resistant and most of the susceptible corn inbreds. The calculated indices reflect species richness (presence or absence of taxa, measured with the ACE and Chao1 indices) and evenness (relative frequencies of taxa); diversity measured with the Shannon and Simpson indices considers both richness and evenness. Specifically, richness differed significantly between resistant and most of the susceptible inbreds for both bacteria and fungi, but only fungal diversity significantly differed over the same inbreds. We also identified the 30 most-abundant bacterial and fungal taxa on these 16 inbreds, and their relative frequencies. Overall, these analyses shed light on the potential impact of *P. maydis* infection on the corn phyllosphere between resistant and susceptible lines.

Our experiment was designed to compare the corn phyllospheric microbiomes between resistant and susceptible inbred lines after natural infection with *P. maydis*. This was necessary because *P. maydis* has not been successfully cultured *in vitro* and appears to be obligately biotrophic. Although the life cycle of *P. maydis* is not known completely, it has been shown to overwinter on corn residue, and spores can be dispersed by rain splash or wind (Groves et al. 2020; Kleczewski et al. 2019; Parbery 1963). The field plot for this experiment is only 14 miles from Lake Michigan and has an excellent environment for the development of tar spot (Telenko et al. 2020). The field plot had been heavily infected with *P. maydis* during the previous year (Telenko et al. 2020), and the conditions yielded a consistent difference in disease severity between resistant and susceptible lines (Table 1). A previous study identified a strong correlation between disease resistance and the phyllosphere microbiome of sorghum (*Sorghum bicolor*) after natural infection (Masenya et al. 2021) in South Africa.

The most abundant bacterial and fungal taxa fell into three likely categories: beneficial, pathogenic, or commensal. Although amplification yielded fewer bacterial reads than fungal, this is not proof that fungal cells outnumbered bacterial cells. Fungi have more rRNA genes per genome, and the fungal and bacterial rDNA fragments were amplified in separate reactions with different primers. Bacterial taxa in both resistant and susceptible lines were dominated by potentially plant-beneficial families *Moraxellaceae*, *Beijerinckiaceae, Erwiniaceae* and *Sphingomonadaceae*, and genera *Methylobacterium*, *Methylorubrum, Pantoea* and *Sphingomonas*. This is consistent with the finding of Wallace et al. (2018) that the vast majority of bacterial reads in corn microbiomes came from only two groups: *Sphingomonads* and the *Methylobacteria*.

Fungal microbial communities were dominated by the families *Phyllachoraceae*, *Cladosporiaceae*, *Didymellaceae* and *Pleosporaceae*, and the genera *Phyllachora*, *Cladosporium*, *Coniothyrium* and *Bipolaris*. *Phyllachora* reads were detected in all inbred lines, even those that were rated the most resistant, and disease severity generally increased with increasing percentage of *Phyllachora* sequences among fungal reads. Exceptions were inbreds CML52, PI685919, PI685950, and PI685788, which were scored as resistant based on disease phenotype with ratings in the range of 0.22 to 3.30 but had very high (65 to 90%) *Phyllachora* sequence reads and thus were considered as susceptible for our analyses.

If the lack of symptoms with relatively high *P. maydis* reads is not a reflection of experimental error, it suggests that stroma development can be suppressed in some corn genotypes or microbial environments that otherwise may permit infection. The most-abundant fungal genera in the resistant lines were *Alternaria, Trichocomaceae* genus*, Ascochyta, Phyllozyma, Papiliotrema*, *Cryptococcus*, *Tilletiopsis, Bipolaris, Stagonospora, Symmetrospora, Epicoccum, Bulleromyces, Coniothyrium* and *Cladosporium*. These genera also predominated in susceptible lines and have been found to be abundant in symptomatic leaves in other studies of tar spot infection (Inacio et al. 2019; McCoy et al. 2019; Singh and Goodwin, 2022; Wagner et al. 2019). Other possible explanations for why some lines showed few visible symptoms but had high *Phyllachora* reads could be that those were latent infections that would show symptoms later, or that the inbreds are tolerant to *P. maydis* so can support a relatively high level of infection with little disease. Additional experiments are required to test these alternative hypotheses.

The composition and frequency of sequence reads provided an indication about other interactions between corn and its associated microbes. For example, although we did not score the inbreds for other diseases, lines that were susceptible could be identified by high numbers of reads for the pathogens that cause southern leaf blight (*Bipolaris maydis*), gray leaf spot (*Cercospora zeae-maydis* or *C. zeina*), and (common or southern) rust (*Puccinia sorghi* or *P. polysora*) (Fig. 5). Distribution of sequence reads also indicated more complex relationships. Reads from *Sphaerellopsis* were rare in inbreds that were resistant to tar spot but were common in three of the more susceptible lines (PI685915, B97, and TX303), the three lines that were clearly susceptible to rust. *Sphaerellopsis filum* is a known mycoparasite of *Puccinia* species (Gordon and Pfender 2012; Trakunyingcharoen et al. 2014) and its frequency most clearly tracked that of its *Puccinia* hosts; it was virtually undetectable in lines that did not also seem to be susceptible to rust (Fig. 2B). Our findings showed that most bacterial and fungal reads were mapped into only a few families and genera, in agreement with a previous study and confirming the relatively low diversity of the corn foliar microbiome (Wallace et al. 2018). While low diversity is consistent with leaf microbiome data from other species (Delmotte et al. 2009), it contrasts sharply with high diversity generally found in the corn rhizosphere (Peiffer et al. 2013; Yang et al. 2017).

The bacterial and fungal alpha diversity metrics showed only a significant difference in microbiota richness (Fig. 3), and fungal community diversity but not bacterial diversity (Fig. 4) between resistant and susceptible lines. Higher species richness (alpha diversity measured by the ACE and Chao1 indices) was observed in most of the resistant lines compared to susceptible inbreds, suggesting that tar spot disease severity (excessive pathogen load) has a negative impact on bacterial and fungal community richness (Fig. 3). This also can be explained by the possibility that beneficial microbes may lose their selective advantage when pathogen load is excessive. Similar to our findings, Southern leaf blight-susceptible plants displayed decreased leaf epiphytic bacterial species richness (Manching et al. 2014).

The finding that only three of 14 susceptible lines did not differ significantly in bacterial and fungal microbiota richness could be explained by the possibility of host genetic effects on these lines (Fig. 3 and Table S1). Previously, it was shown that the genetic composition of the host structured the phyllosphere microbiome (da Silva et al. 2016; Wagner et al. 2019; Wallace et al. 2018). However, we did not see a clear difference in bacterial community diversities between the resistant and susceptible lines (Fig. 4). Except for the susceptible inbred lines PI685790 and TX303, which were significantly lower, the other fourteen lines showed similar levels of bacterial community diversities (Fig. 4, A and B). However, there was a significant difference in fungal community diversity between resistant and susceptible lines depending on the sample; the twelve most-susceptible lines (CML52, PI685915, PI685790, PI685919, PI685950, PI685836, PI685918, B97, 4401350, PI685788, TX303, and PI685920) had significantly lower fungal community diversities compared to those of the resistant lines CML103 and CML69 (Fig. 4, C and D). These findings suggest that *P. maydis* infection has a significant impact primarily on the richness of the phyllosphere microbiome and its fungal diversity but not bacterial diversity when including evenness. Nevertheless, we cannot ignore the likelihood of host genetic influences on community diversity. In a previous study, corn recombinant inbred lines (RILs) revealed a major link between plant genetics, community diversity, and disease resistance (Balint-Kurti et al. 2010). Furthermore, high bacterial diversity in maize lines was linked to SLB susceptibility in the field (Balint-Kurti et al. 2010). Our findings contrast with those of Wallace et al. (2018), who found that corn genotype did not affect diversity of the foliar microbial community in an RIL population. Possibly the genetic diversity in the population analyzed by Wallace et al. (2018) was lower than those in our panel of inbreds or the RIL population analyzed by Balint-Kurti et al. (2010). It is also possible that environmental heterogeneity was great enough to obscure any genetic effects in the Wallace et al. (2018) study.

Two measures of interpopulation similarity as measured by the beta-diversity indices demonstrated highly significant heterogeneity of microbial populations among inbreds, with *p-* values less than 2e^-16^ after ANOVA of the Sorensen and Bray-Curtis metrics. These two indices compare, respectively, taxon presence or absence and the relative populations of mostly the same taxa among inbreds. Because the field environment was macroscopically if not temporally uniform, this qualitative and quantitative heterogeneity is most likely determined by the genotype of each inbred, which sets the stage for secondary interactions among microbial taxa and interactions with host defense responses.

Bacterial genera *Nocardioides, Geodermatophilus, Kineosporiaceae* genus, *Angustibacter, Methylorubrum*, and *Quadrisphaera* were more abundant in resistant than susceptible inbreds (Fig. 5A). These beneficial genera promote stress tolerance and potentially improve overall plant growth, as illustrated by the following examples. *Nocardioides thermolilacinus*, a potential biocontrol agent in tomato, reduces disease severity due to foliar fungal pathogens *Alternaria solani* and *Corynespora cassiicola*, the oomycete *Phytophthora infestans*, and the bacterial pathogens *Pseudomonas syringae* pv. *tomato* and *Xanthomonas campestris* pv. *vesicatoria*, in both laboratory and field trials (Filho et al. 2008). *Methylorubrum extorquens* benefits host development and stress tolerance by altering plant hormone signaling pathways, downregulating senescence and cell death-associated genes, and inducing ononitol biosynthesis in pine trees (Koskimäki et al. 2022). Isolates of *Methylorubrum*, *Marmoricola*, *Geodermatophilus* and *Nocardioides* promoted stress tolerance and exhibited antimicrobial activity against a variety of pathogens (Jiang et al. 2018; Sun et al. 2015) and may be of interest for controlling tar spot of corn.

In contrast, some bacteria promote plant disease directly as pathogens or indirectly by enhancing susceptibility to other pathogens. *Spirosoma, Aureimonas, Deinococcus, Pseudomonas*, and *Pantoea* were elevated in the susceptible lines (Fig. 5A). *Pantoea*, *Enterobacter* and *Pseudomonas* include opportunistic pathogens that have been reported to cause disease in corn (Hinton and Bacon 1995; Roper 2011; Vidaver and Carlson 1978). Abundant genera *Methylobacterium*, *Sphingomonas* and *Klenkia* were associated with both resistant and susceptible lines, and they have been frequently observed in the phyllosphere of corn.

Fungal genera *Cladosporium*, *Tilletiopsis*, *Cryptococcus* and *Papiliotrema* were associated with the resistant lines (Fig. 5B). Members of these four genera have been demonstrated to be quite efficient at suppressing various plant pests and diseases. For example, a *Cladosporium* species suppressed whiteflies and aphids in a greenhouse (Abdel-Baky and Abdel-Salam 2003), *Tilletiopsis* spp. have inhibited powdery mildews (Kiss 2003; Last 1970), *Cryptococcus laurentii* has suppressed blue mold and brown rot in sweet cherry (Qin and Tian 2004), and *Papiliotrema* spp. have controlled postharvest pathogens (Castoria et al. 2021). These findings suggest that these fungal genera may be excellent candidates for further investigation into tar spot disease prevention. On the other hand, in addition to *Phyllachora* itself, *Sphaerellopsis*, *Puccinia*, and *Didymella* increased relatively in the susceptible inbreds, suggesting that these taxa may have contributed to tar spot susceptibility or disease progression (Fig. 5B).

Potentially beneficial bacterial and fungal taxa declined in susceptible lines in response to increased *P. maydis* load, implying that tar spot severity affects the bacterial and fungal communities. As the disease progresses, *P. maydis* might displace or inhibit these beneficial microbial taxa for its own advantage, further aggravating the disease. Previously, Manching et al. (2014) demonstrated that disease status shapes phyllospheric microbiome composition and structure. Meanwhile, other plant-pathogenic bacterial and fungal taxa increased in susceptible lines with increased *P. maydis* reads, indicating that these microbes are associated with tar spot disease progression.

In general, host genotype, weather, and availability of inoculum limit variation in the foliar microbiome. At least six specific relationships can operate within these limits:

1. *Phyllachora* invasion might physically displace a pre-existing microflora without changing its composition. In this case, reduced richness would reflect decreasing sample size of non-*Phyllachora* reads.
2. *Phyllachora* competes with a co-existing microflora for water and nutrients from host cells, and the less-proficient competitors die out.
3. *Phyllachora* could produce soluble compounds, for example antibiotics, that directly antagonize other taxa.
4. *Phyllachora* triggers host defense responses that differentially affect other organisms.
5. The corn inbreds vary in genotype for receptors that allow *Phyllachora* to detect and attack host cells. Possibly these receptors also differentially affect other taxa.
6. Other organisms could chemically antagonize *Phyllachora*, and their antibacterials or antifungals could collaterally inhibit other microbes.

While physical displacement obviously occurs, our data suggest that one or more of the other mechanisms also operate in tar spot disease. To the best of our knowledge, this is the first analysis to investigate the bacterial and fungal microbiomes in the phyllospere of corn in relation to natural infection with *P. maydis*. We showed that this infection results in distinct phyllospheric microbial communities in resistant and susceptible corn inbreds. Taxa that are differentially abundant between resistant and susceptible lines might be of interest for their potential for controlling tar spot of corn. Organisms with strong positive correlations with *Phyllachora* reads could be possible mycoparasites, which increase in abundance with greater availability of their food supply. The strong association of the known rust mycoparasite *S. filum* with reads of its *Puccinia* host supports this idea. Conversely, organisms that are highly negatively correlated with *Phyllachora* reads could be antagonists that compete directly or produce compounds that inhibit tar spot development. Either group could be a source of organisms for possible biocontrol of *P. maydis*. Functional characterization of these beneficial, differentially abundant taxa might elucidate important details of their interactions with *P. maydis* that indicate how they could be deployed to mitigate tar spot disease in the future.

### Data availability

Raw reads data have been uploaded and processed by NCBI. The bioproject number is PRJNA1044151.

## ACKNOWLEDGMENTS

We thank Dr. Tiffany Jamann at the University of Illinois, Urbana-Champaign and Dr. Gurmukh Johal at Purdue University, for providing seeds of the GEM and NAM population parents, respectively, and Jeffrey Ravellette at Purdue University for assistance with field trial establishment and maintenance.

## Supplemental Figures

**Supplementary Fig. S1.** The 30 most abundant bacterial phyla in 16 inbred corn lines sorted from left to right as resistant or susceptible based on the number of *Phyllachora maydis* sequence reads. The values represent the relative abundances averaged over three replications. The color matrix indicates the designated taxonomic unit.

**Supplementary Fig. S2.** The 30 most abundant fungal phyla in 16 inbred corn lines sorted from left to right as resistant or susceptible based on the number of *Phyllachora maydis* sequence reads. The values represent the relative abundances averaged over three replications. The color matrix indicates the designated taxonomic unit.

**Supplementary Table S1.** Tests for significance of differences between measures of alpha diversity between resistant and susceptible corn lines determined based on the numbers of reads to the fungal pathogen *Phyllachora maydis* among 16 inbred lines chosen from the parents of the NAM and GEM populations. Significant differences were calculated by one-way ANOVA with R function “aov”, and pairwise estimates of significance were calculated with Tukey’s Honest Significant Difference test (R function “TukeyHSD”) (*p* ≤ 0.05).

