## Supplemental Figure 2 for "Tar Spot Disease Severity Influences Phyllosphere-Associated Bacterial and Fungal Microbiomes"

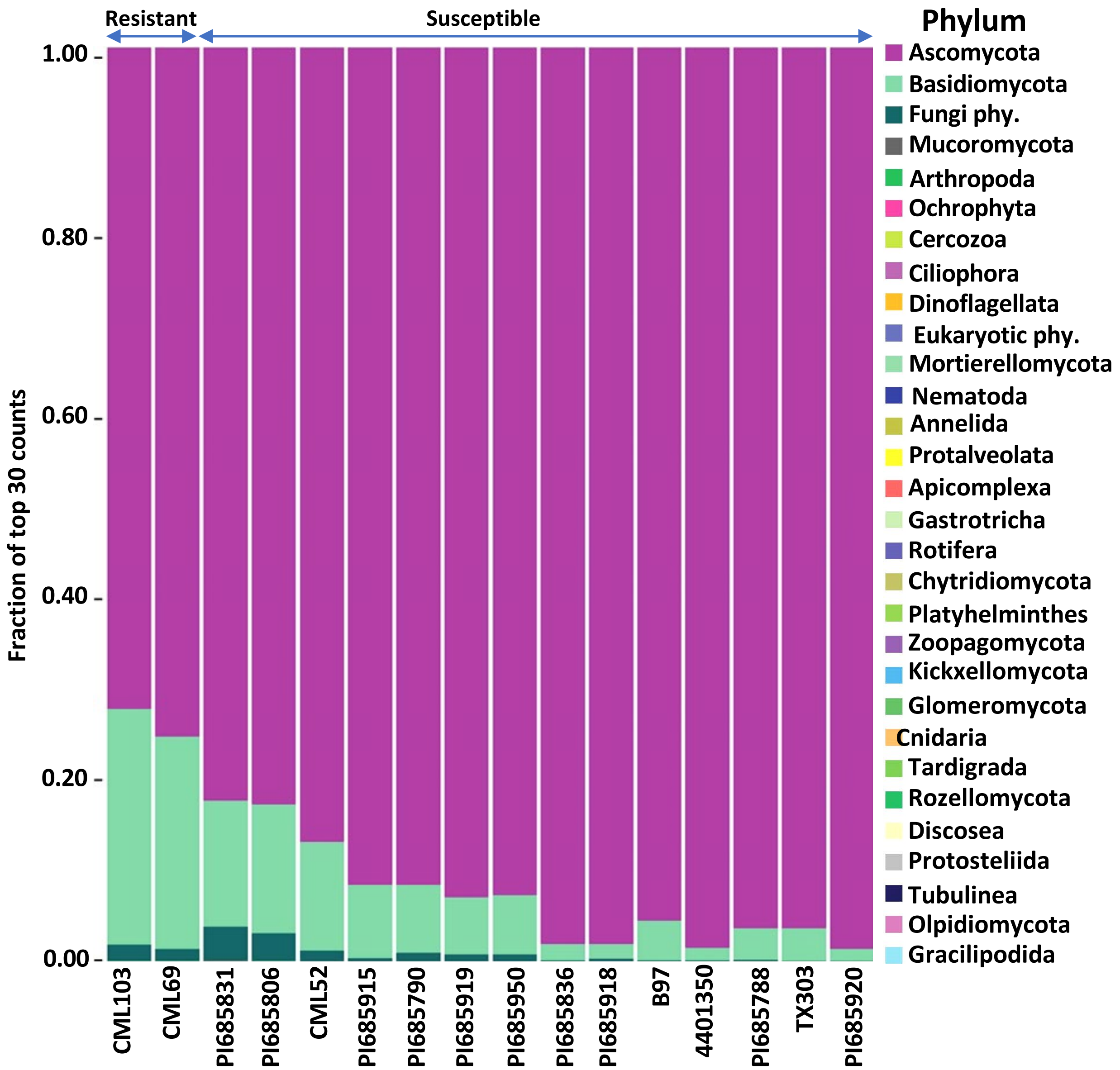


Fig. S2. The 30 most abundant fungal phyla in *Phyllachora maydis* resistant and susceptible corn lines.
The values represent the mean relative abundances (n=3). The color matrix indicates the designated taxonomic unit.
